# Influenza A Virus NS1 Limits Recognition of Double-Stranded Transposable Elements by Cytosolic RNA Sensors

**DOI:** 10.1101/2024.05.24.595739

**Authors:** Marie Lork, Liam Childs, Gauthier Lieber, Renate König, Benjamin G. Hale

**Affiliations:** Institute of Medical Virology, University of Zurich, 8057 Zurich, Switzerland; Host-Pathogen Interactions, Paul-Ehrlich-Institut, 63225 Langen, Germany

**Keywords:** dsRNA sensors, influenza A virus, transposable elements, viral antagonism

## Abstract

Influenza A virus (IAV) infection triggers de-repression of host transposable elements (TEs), which have the potential to form double-stranded (ds)RNAs and stimulate innate antiviral immunity. However, as wild-type IAV is generally a poor inducer of innate immunity, it remains unclear whether de-repressed TEs actually form dsRNAs recognizable by host cytosolic RNA sensors, or whether IAV might antagonize such sensing. Here, we performed strand-specific total RNA-Seq on nuclear and cytosolic fractions from cells infected with wild-type IAV or a recombinant IAV lacking NS1, a viral dsRNA-binding protein. Both infections led to global increases in host TE RNAs with bioinformatic and experimental evidence for double-strandedness. However, only NS1-deficient IAV infection led to significant amounts of TE-dsRNAs translocating to the cytosol, and co-precipitations identified that wild-type NS1 associates with TE-dsRNAs. Furthermore, a functional screen indicated that TE-dsRNAs can be engaged by various host cytosolic RNA sensors, including RIG-I, MDA5, ZBP1, and PKR. Our data reveal the double-stranded nature of infection-triggered host TEs and suggest an NS1-mediated sequestration mechanism to limit their cytosolic abundance and broad activation of diverse sensors.

## Introduction

While only 1-2% of the human genome encodes proteins, the remaining repetitive, low-complexity part is poorly characterized, and was long referred to as ‘junk DNA’ (ENCODE Project Consortium, 2012; Lander *et al*, 2001). A substantial fraction thereof (∼45% of the human genome) is made up by transposable elements (TEs), DNA sequences that have the ability to move within a genome and can be classified based on their mechanism of transposition: DNA transposons move by a ‘cut and paste’ mechanism while retrotransposons adopt a ‘copy and paste’ mechanism using their RNA as an intermediate (Luning Prak & Kazazian, 2000). Endogenous retroviruses (ERVs), a type of retrotransposon, stem from ancient retroviral insertions into germline cells and feature two long terminal repeats (LTRs) flanking their proviral genome (Grundy *et al*, 2021). Aside from LTR-containing ERVs, the genome also harbors non-LTR retroelements, like long interspersed elements (LINE1s) and short interspersed elements (SINEs). TE retrotransposition is generally associated with genome instability due to target gene disruption, alternative splicing, and various epigenetic changes (Bhat *et al*, 2022). For example, it has been demonstrated that the global epigenetic dysregulation of TEs in cancer cells, resulting in their expression, can have oncogenic effects in certain scenarios (Grundy *et al*., 2021). To avoid these undesirable effects, expression of TEs is usually repressed by epigenetic mechanisms such as DNA methylation and histone modification. One key silencing pathway in mammals involves Krüppel-associated box (KRAB) zinc-finger proteins (ZFPs), which bind to specific DNA sequences in LTR elements and, together with TRIM28 (KAP1), recruit the methyltransferase SETDB1 to the promoter regions of KRAB target genes resulting in repression of specific TEs (Rowe *et al*, 2010; Turelli *et al*, 2014). More recently, the involvement of the Human Silencing Hub (HUSH) complex in the suppression of TEs, especially evolutionary young LINE1 elements, has been described (Robbez-Masson *et al*, 2018).

Intriguingly, TEs have gained considerable interest for their apparent co-option by cells as contributors to innate immune defenses (Hale, 2022). For example, TE LTRs can function as promoters or enhancers for nearby genes, thereby affecting gene regulation, including for genes involved in the innate immune response (Chuong *et al*, 2016; Srinivasachar Badarinarayan & Sauter, 2021). Additionally, aberrant accumulation of de-repressed RNA transcripts can be a source of highly immunogenic ‘self’ dsRNA, that can be sensed by host pattern recognition receptors (PRRs) to induce an antiviral interferon (IFN) response (Chiappinelli *et al*, 2015; Cuellar *et al*, 2017; Roulois *et al*, 2015). Mechanistically, the production of immunostimulatory TE-dsRNAs can arise through various processes, including LTR-dependent bidirectional transcription, hairpin formation, or read-through transcription of head-to-head or tail-to-tail arranged elements (Sadeq *et al*, 2021). In this context, we recently identified a new type of physiological mechanism leading to activation of antiviral immunity in response to virus infection: an influenza A virus (IAV)-triggered ‘switch’ in TRIM28 causes transcriptional de-repression of host TE RNAs (mainly ERVs) and the initiation of a protective IFN response (Schmidt *et al*, 2019). Specifically, we could show that loss of SUMO-modified TRIM28 leads to the transcriptional upregulation of several TEs that may act in an immunostimulatory fashion via the PRR, retinoic acid-inducible gene-I (RIG-I), to impair virus replication. These data not only indicated that TEs have been co-opted to aid defense against exogenous pathogens, but that the mechanisms governing retroelement silencing are physiologically regulated by the cell. Notably, IAV is not the only type of pathogen described to upregulate various host retroelements during infection (Young *et al*, 2014). Recently published re-analyses of publicly available gene expression datasets showed that infection with a diverse range of viruses can trigger changes in TE expression (Chen *et al*, 2023; Macchietto *et al*, 2020), indicating that the insights on molecular mechanisms leading to IAV-induced TE expression and sensing could be broadly applicable to other viruses.

To date, it has only been suggested that IAV-triggered de-repressed retroelements have the potential to promote antiviral immunity by forming immunostimulatory dsRNAs that might activate the cytosolic PRR, RIG-I (Schmidt *et al*., 2019). Indeed, it was not formally shown that these TEs form dsRNA, or that they can exit the nucleus and be located in the cytosol where several PRRs, including RIG-I and many more, are based. In addition, it is still unclear whether IAV might antagonize this pathway downstream of TE de-repression, as wild-type (wt) IAV is known to employ various countermeasures to limit host antiviral defenses (Hale *et al*, 2010). In this context, the non-structural protein 1 (NS1), is probably the most well-characterized of the IAV innate immune antagonist proteins (Ayllon & García-Sastre, 2015). NS1 contains a dsRNA-binding domain that is critical for suppressing IFN induction during IAV infection (Donelan *et al*, 2003), although the precise dsRNA target bound by NS1 remains unclear (Cheng *et al*, 2009; Hatada *et al*, 1992; Weber *et al*, 2006; Zhang *et al*, 2018).

To address these open questions, we performed an in-depth transcriptomic analysis of nuclear and cytosolic fractions obtained from cells infected with wt IAV or a recombinant IAV lacking NS1 expression (ΔNS1). We confirmed that lack of NS1 leads to enhanced stimulation of host innate immune responses during infection, as characterized by the expression of cytokines, chemokines, and interferon-stimulated genes (ISGs). Employing a novel bioinformatics analysis pipeline, we further identified and quantified the expression of TEs in each cellular compartment during infection, focusing on TEs that exhibited overlapping sense and anti-sense transcripts and thus having the potential to form dsRNA. We observed a significant increase in such potentially double-stranded TE RNAs (TE-dsRNAs) during infection with both viruses, an observation that could be validated orthogonally using an antibody specific to dsRNA. Strikingly, our fractionation analysis revealed that TE-dsRNAs were predominantly localized in the cytosol only following infection with the ΔNS1 virus, suggesting a role for NS1 in normally restricting TE-dsRNA exit from the nucleus. This hypothesis was supported by the finding that wt NS1, but not a dsRNA-binding deficient mutant, could co-precipitate TE-dsRNAs. Finally, we found that, in the absence of NS1, infection-triggered TE-dsRNAs can be engaged by a range of cytosolic dsRNA PRRs, such as RIG-I, MDA5, ZBP1, and PKR, indicating that failure of NS1 to antagonize TE-dsRNA function may broadly impact immunostimulatory, inflammatory and cell death antiviral pathways.

## Results

### Integrated Analysis of Subcellular Host Transcriptomes During IAV Infection Reveals the Importance of NS1 in Suppressing Cytokine and Chemokine Induction

To gain comprehensive information on expression of protein coding genes and TEs during IAV infection, we established an in-depth transcriptomic analysis workflow (**Fig 1A**). Given the importance of IAV NS1 as a known IFN-antagonist (Ayllon & García-Sastre, 2015), we sought to compare the differential transcriptional responses induced by wt IAV or IAV ΔNS1 (both based on A/WSN/1933, WSN; H1N1). We therefore performed host RNA sequencing of an A549-based human lung epithelial cell line at 8 or 16 hours post infection (hpi) with either wt IAV or IAV ΔNS1 at an MOI of 5 PFU/cell, and compared this with mock infected cells. Importantly, following initial cell harvest, we separated cells into nuclear and cytosolic fractions, and extracted total RNA from these fractions to allow an understanding of the subcellular localization of examined transcripts. To enhance resolution and simplify sample complexity, ribosomal RNA depletion was performed prior to sequencing. We opted for ribosomal depletion over the polyadenylation (polyA) enrichment method to ensure comprehensive transcriptomic coverage, mitigating the risk of overlooking TE transcripts lacking a polyA tail. Furthermore, the experiment and subsequent bioinformatics analysis was specifically designed to evaluate the presence of de-repressed TEs in a double-stranded state: RNA sequencing was conducted in a strand-specific manner, thus facilitating subsequent assignment of reads to their original strands, and downstream bioinformatics analysis focused on determining relative changes in TE transcripts with overlapping sense and antisense transcripts.

**Figure 1:**
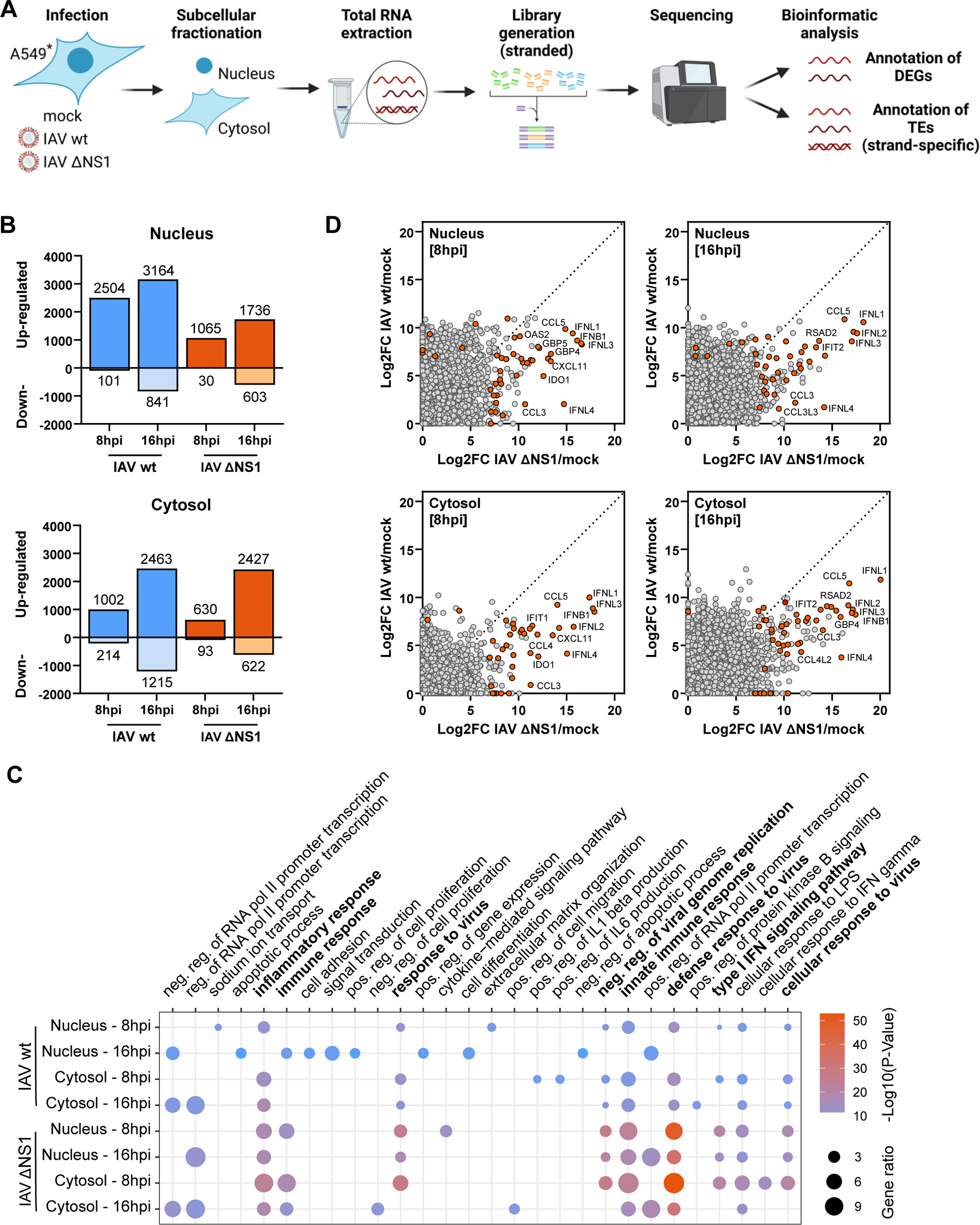
Integrated Analysis of Subcellular Host Transcriptomes During Infection Reveals the Importance of NS1 in Suppressing IAV-induced Cytokines and Chemokines. **A.** Schematic representation of the experimental set-up: illustration depicting the experimental design, whereby A549 cells (*A549-ACE2/TMPRSS2) were infected with wt IAV or IAV ΔNS1 [MOI 5 PFU/ml] for 8h or 16h, followed by subcellular fractionation, RNA extraction, and transcriptome analysis. For sequencing analysis n=3 independent replicates were performed. **B.** Transcriptome analysis of subcellular fractions: comparison of the total number of differentially expressed genes (DEGs) in distinct subcellular fractions of A549-ACE2/TMPRSS2 cells following infection with wt IAV or IAV ΔNS1 [MOI 5 PFU/ml] at the times indicated. DEGs were defined by log_2_FC > 2 and Padj < 0.1. **C.** Gene ontology analysis with DAVID: gene ontology (GO) analysis using DAVID was performed on DEGs with log_2_FC > 2 and Padj < 0.1. Dot plot visualization displays the top 10 enriched GO terms for each condition (P < 0.001). Dot color indicates the P-value, while dot size represents the percentage of genes enriched in the total gene set. **D.** Comparative analysis of DEGs: comparison of DEGs between wt IAV and IAV ΔNS1-infected subcellular fractions at the indicated times. Cytokines, chemokines, and interferon-stimulated genes (ISGs) with a log_2_FC > 7 in either condition are highlighted in orange, emphasizing genes with substantial expression changes.

In all replicates of our sequencing experiment, verification of subcellular fraction purity was achieved using RT-qPCR for established markers: the long non-coding RNA MALAT1 for nuclear localization (Hutchinson *et al*, 2007), and GAPDH mRNA for its increased prevalence in the cytosol (Samacoits *et al*, 2018) (**Fig EV1A**). Additionally, western blot analysis for Histone H3 served as a nuclear marker and for GAPDH as a cytosolic marker (**Fig EV1B**). Transcriptomic profiling of the fractionated samples revealed substantial alterations in cellular gene expression following both IAV infections compared to mock-infected cells across all timepoints and fractions (**Fig 1B**, and **Dataset EV1**). Infection with wt IAV for 8 or 16 hours resulted in 2504 and 3164 genes, respectively, exhibiting significant induction (log_2_ fold change >2; adjusted p-value <0.1) in the nuclear fraction. Additionally, 1002 and 2463 genes, respectively, were upregulated in the cytosolic fraction upon infection with wt IAV (**Fig 1B**). Infection with IAV ΔNS1 resulted in comparable numbers of differentially expressed genes (DEGs) in the cytosolic fraction as compared to wt IAV infection, with 630 and 2427 genes significantly upregulated after 8 or 16 hours, respectively (**Fig 1B**). This was also broadly the case in the nuclear fraction, with IAV ΔNS1 infection leading to about half the number of DEGs as compared to wt IAV infection. Gene enrichment analyses of DEGs revealed that while both wt IAV and IAV ΔNS1 modulated similar pathways such as ‘response to virus’, ‘innate immune response’, and ‘inflammatory response’, IAV ΔNS1 exhibited much stronger effects (**Fig 1C**). This observation was further substantiated through direct comparison of specific gene expression changes between wt IAV and IAV ΔNS1-infected cells: both virus infections elicited upregulation of RNAs encoding cytokines, chemokines, and interferon-stimulated genes (ISGs) across both subcellular fractions, but the magnitude of induction was notably greater in IAV ΔNS1-infected cells (**Figs 1D**, **EV1C**, and **Dataset EV1**). To rule out that these effects were caused by potential differences in viral replication kinetics, we used RT-qPCR to confirm similar levels of viral M segment RNA between wt IAV and IAV ΔNS1 infections (**Fig EV1D**). Taken together, these transcriptomic analyses reveal distinct patterns of host DEGs during infection of A549-based cells with wt IAV and IAV ΔNS1, and highlight the role of NS1 in limiting induction of key immune and inflammatory pathways.

### Bioinformatics Analysis of IAV-Induced Double-Stranded TEs Identifies Increased Cytosolic Abundance in the Absence of NS1

We next sought to comprehensively characterize TE upregulation during IAV infection and assess the evidence for the production of double-stranded TE RNAs (TE-dsRNAs). For this purpose, we developed a bioinformatics pipeline for the *in silico* detection of dsRNA from stranded, paired, short read RNA-seq data. The pipeline initially trimmed and aligned reads to the human genome allowing gapped alignments. It then counted reads that both mapped uniquely to TEs and had the potential to form dsRNA with reads from the opposing strand, yielding estimates of dsRNA read counts for each TE. Finally, the differential expression of TE-dsRNA transcripts was calculated. An illustrative depiction of the annotation of sense and antisense reads is presented in **Fig 2A**. The bioinformatics analysis unveiled a substantial upregulation of TEs with reads in both sense and antisense orientations in the nucleus in response to infection with both wt IAV and IAV ΔNS1, which is suggestive of sequences that could form potential TE-dsRNAs (**Figs 2B-C**, **EV2A**, and **Dataset EV2**). Specifically, cells infected with wt IAV for 8 or 16 hours exhibited significant upregulation of 4365 and 6329 TE-dsRNAs (log_2_ fold change >2; adjusted p-value <0.1), respectively, in the nuclear fraction compared to mock-infected cells (**Fig 2B**). Remarkably, only a low number of these TE-dsRNAs were detected in the cytosolic fraction during wt IAV infection (83 and 196 at 8 hpi and 16 hpi, respectively) (**Fig 2B**). Conversely, infection with IAV ΔNS1 led to fewer differentially expressed TE-dsRNAs in the nucleus, with only 1415 and 1656 TE-dsRNAs upregulated after 8 h and 16 h of infection, respectively (**Fig 2B**). However, it was striking that cells infected with IAV ΔNS1 exhibited a substantially greater number of potential TE-dsRNAs in the cytosol (190 and 754 at 8 hpi and 16 hpi, respectively) (**Fig 2B**). Collectively, the proportion of cytosolic versus nuclear TE-dsRNAs was markedly higher in IAV ΔNS1-infected cells compared to wt IAV-infected cells (**Figs 2C-D**). Specifically, at 16 h post IAV ΔNS1 infection, about 30% of the total TE-dsRNAs were localized in the cytosol, whereas in wt IAV-infected cells, only 3% of the induced TE-dsRNAs demonstrated cytosolic localization (**Fig 2D**). Notably, this enhanced cytosolic localization of TE-dsRNAs in IAV ΔNS1 infected cells appeared to be specific to this class of RNAs, as no clear differences in the nuclear and cytosolic distribution of non-TE DEGs were observed between wt IAV and IAV ΔNS1 (**Fig 2E**). Furthermore, multiple different TE families and classes were observed as dsRNAs in the cytosol, including LTRs, LINEs, SINEs, and DNA transposons, and we did not observe any family being notably overrepresented or any potential length bias, e.g. towards the shorter Alu/SINE elements (**Fig EV2B-C**). These sequencing data provide evidence that IAV infection not only leads to substantial induction of TEs (Schmidt *et al*., 2019), but that many of these TEs have the potential to form TE-dsRNAs via sequence complementarity. Furthermore, these fractionation data reveal that such TE-dsRNAs can reach the cytosol of IAV-infected cells, particularly in the absence of the IAV NS1 protein.

**Figure 2:**
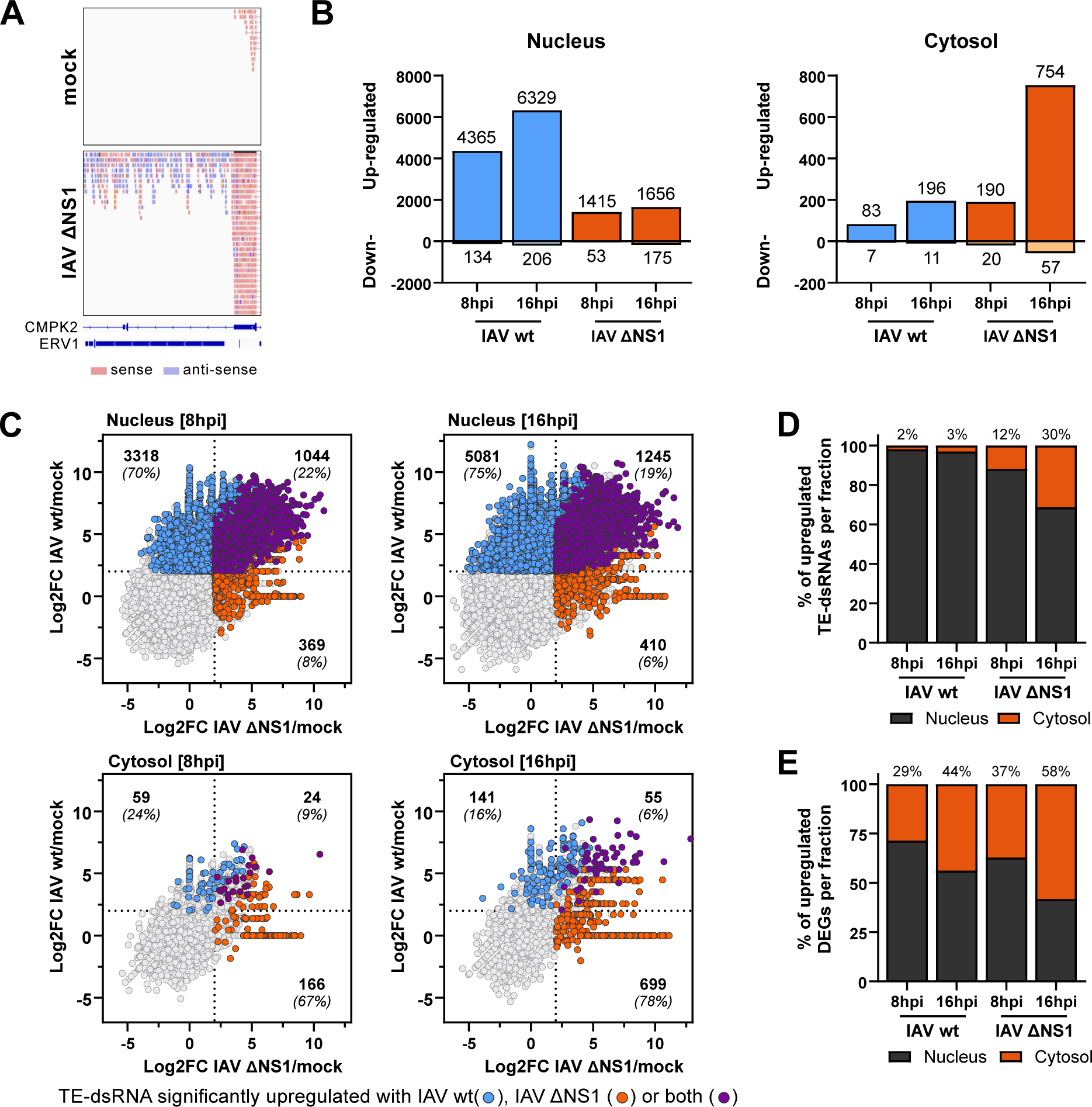
IAV Infection Leads to Upregulation of Double-Stranded TEs, which Translocate to the Cytosol in the Absence of NS1. **A.** Bioinformatic strategy for read filtering: in our bioinformatic approach, reads expressed in both directions were filtered for and assessed as likely to form double-stranded RNA (dsRNA). Forward reads are shown in red, while reverse reads are depicted in blue. **B.** Transcriptome analysis of differentially expressed transposable elements (TEs) in subcellular fractions: quantification of differentially expressed TEs (DE-TEs) with evidence for double-strandedness (log_2_FC > 2; Padj < 0.1) in nuclear and cytosolic fractions following infection with wt IAV or IAV ΔNS1 [MOI 5 PFU/ml] for the indicated times. **C.** Comparative analysis of double-stranded DE-TEs in response to wt IAV or IAV ΔNS1 infection: comparative visualization of significantly upregulated double-stranded DE-TEs in response to wt IAV (blue) and IAV ΔNS1 (orange) infections. TEs significantly upregulated in both conditions are highlighted in purple. The numbers in each quadrant represent both the absolute counts and the relative percentages of double-stranded DE-TEs that were significantly upregulated. **D.** Percentage of double-stranded DE-TEs in distinct subcellular fractions: bar chart of the percentage of double-stranded DE-TEs (log_2_FC > 2; Padj < 0.1) within each respective subcellular fraction during wt IAV and IAV ΔNS1 [MOI 5 PFU/cell] infections at the indicated timepoint. The percentages annotated above the bars specifically denote the cytosolic fractions. **E.** Percentage of DEGs in distinct subcellular fractions: bar chart of the percentage of DEGs (log_2_FC > 2; Padj < 0.1) within the respective subcellular fractions at the indicated times post infection. The percentages annotated above the bars specifically denote the cytosolic fractions.

### dsRNA-Immunoprecipitation Assays Validate the Double-Stranded Nature of IAV-Induced TEs and their Increased Cytosolic Abundance in the Absence of NS1

To provide orthogonal evidence that IAV infection leads to upregulation of TE-dsRNA species, we established a protocol to immunoprecipitate dsRNA from cells and quantify specific TE RNA transcripts by RT-qPCR (**Fig 3A**). Specifically, A549 cells were infected with either wt IAV or IAV ΔNS1 for 16 hours, after which total RNA was harvested using Trizol extraction, and dsRNA species were immunoprecipitated using the well-characterized anti-dsRNA antibody, 9D5 (Son *et al*, 2015). RNA was then extracted from the resulting immunoprecipitates using Trizol and was subsequently analyzed by RT-qPCR for a selection of TEs for which we had transcriptomic and bioinformatic evidence for double-strandedness (LTR13, MER72, MER21B, L1MD2, MLT1F, MLT2B3). All TEs examined were clearly enriched in immunoprecipitates from the 9D5 antibody as compared to the control antibody (anti-Flag) (**Fig 3B**). Furthermore, TE-dsRNA was detected in response to infection with both viruses, wt IAV as well as IAV ΔNS1. These data support the findings from our RNA sequencing and bioinformatics analyses, and further indicate that these IAV-induced TEs can form dsRNA species.

**Figure 3:**
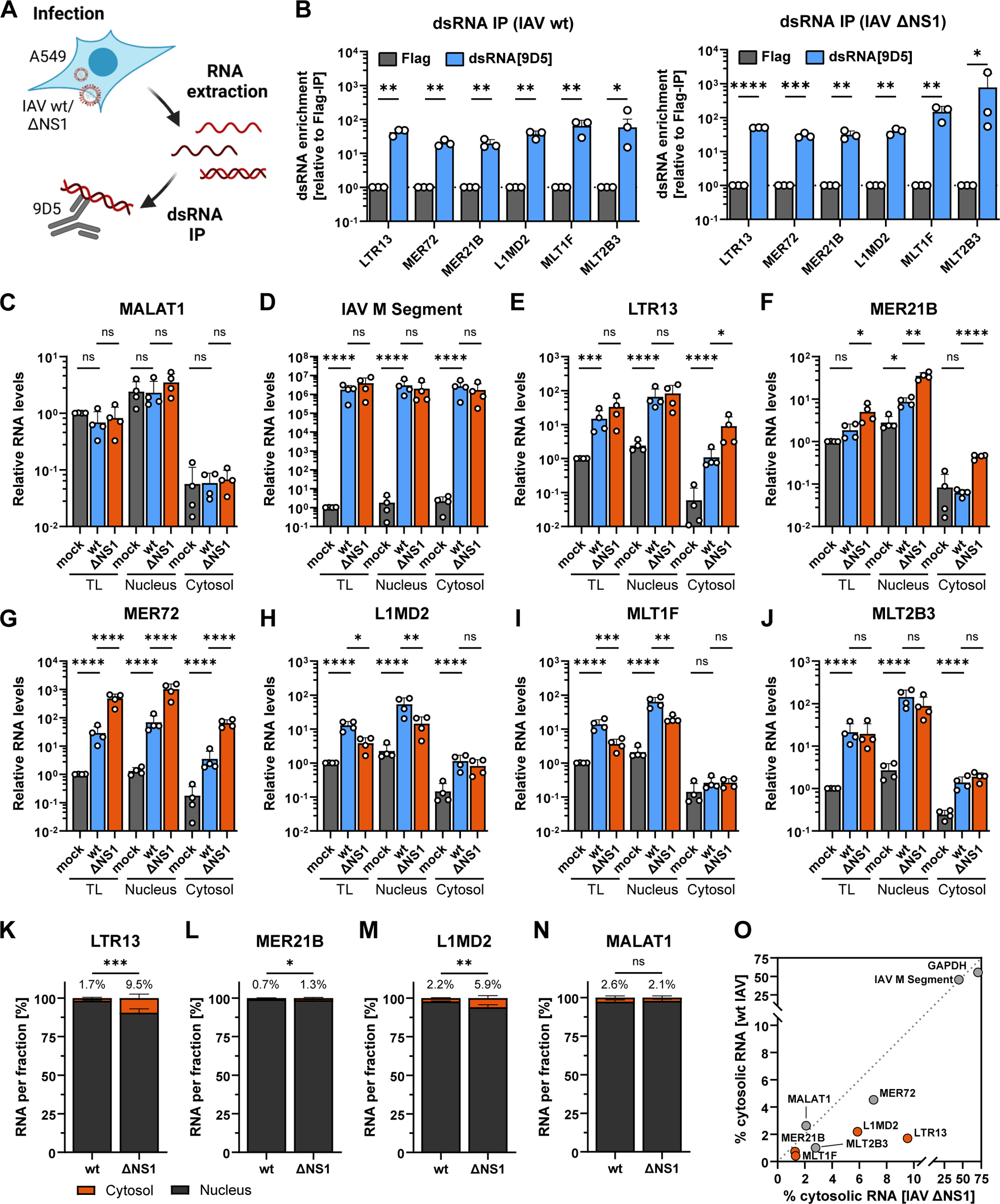
dsRNA-Immunoprecipitation Assays Validate the Double-Stranded Nature of IAV-Induced TEs and their Increased Cytosolic Abundance in the Absence of NS1. **A.** Experimental set-up for dsRNA immunoprecipitations: schematic representation of the experimental design aimed at discerning the double-stranded nature of upregulated TEs by immunoprecipitation (IP) with a specific dsRNA antibody (clone 9D5) from total extracted RNA of IAV-infected cells. **B.** dsRNA immunoprecipitation: A549 cells were infected with wt IAV or IAV ΔNS1 [MOI 5 PFU/ml] for 16 hours, before total RNA was extracted and dsRNA IP was performed using a specific dsRNA antibody (9D5). Selected TEs were subsequently analyzed via RT-qPCR. Enrichment is presented as fold change over the negative control (IP with anti-Flag antibody). Bars represent mean values and SDs from n=3 independent experiments (each dot corresponds to one replicate). Statistical significance was determined by one-sample t-test on log-transformed data, comparing the sample means to their individual controls (*P ≤ 0.05; **P ≤ 0.01; ***P ≤ 0.001; ****P ≤ 0.0001). **C-J.** Validation of TE expression and localization by RT-qPCR: validation of differential expression of selected TEs in subcellular fractions following mock, wt IAV, or IAV ΔNS1 infections [MOI 5 PFU/ml; 16 hpi] in A549 cells. MALAT1 served as a nuclear localized fractionation control. Bars represent mean values and SDs from n=4 independent experiments (each dot corresponds to one replicate). Significance was determined by ordinary one-way ANOVA with Šídák’s multiple comparisons test on log-transformed data (*P ≤ 0.05; **P ≤ 0.01; ****P ≤ 0.0001; ns, non-significant). **K-N.** Expression ratios of selected TEs: quantification of selected TEs determined via RT-qPCR, depicted as the percentage of transcript per fraction. Bars represent mean values and SDs from n=4 independent experiments. Significance was determined by unpaired t test (*P ≤ 0.05; **P ≤ 0.01; ***P ≤ 0.001; ns, non-significant). **O.** Cytosolic presence of TEs between wt IAV and IAV ΔNS1-infected cells: comparison of the cytosolic presence of selected TEs and other genes between wt IAV and IAV ΔNS1-infected cells. The percentage of each TE present in the cytosolic fraction (versus the nuclear fraction) of each infected condition is plotted in a YX graph. Each dot represents the mean of the four independent experiments shown in C-J. Transcripts with a statistically significant cytosolic localization upon IAV ΔNS1 infection as compared to wt IAV infection are highlighted in orange (data from panel K-N and Fig EV3). GAPDH and IAV M segment RNAs were used as controls for predominantly cytosolic localizations.

We next specifically assessed the IAV-induced subcellular distribution of TEs for which we had both RNA sequencing and dsRNA-immunoprecipitation data supporting their double-strandedness. In particular, we compared the impact of wt IAV and IAV ΔNS1 infection on TE-dsRNA localization. A549 cells were infected with wt IAV or IAV ΔNS1 at an MOI of 5 PFU/cell for 16 hours (or mock), and RT-qPCR for specific candidate TE-dsRNAs was subsequently performed on RNA extracted from total cell lysates or from nuclear and cytosolic fractions. RT-qPCR for the nuclear marker MALAT1 confirmed successful fractionation, and that there were no gross differences in nuclear/cytosolic localization of this RNA species between wt IAV and IAV ΔNS1 infected cells (**Fig 3C**). In addition, RT-qPCR for the IAV M segment confirmed similar infection levels between wt and ΔNS1 viruses (**Fig 3D**). Furthermore, all the selected TEs assayed were clearly induced by both wt IAV and IAV ΔNS1 and were found in both the nuclear and cytosolic fractions (**Figs 3E-J**). However, when comparing infections with wt IAV and IAV ΔNS1, it was clear that IAV ΔNS1 infection resulted in significantly higher amounts of cytosolic RNA for many TEs (**Figs 3E-J**), which was particularly evident when the data were expressed as the ratio between cytosolic and nuclear transcript abundance (**Fig 3K-M, O, Fig EV3**). Such increased cytosolic localization upon IAV ΔNS1 infection as compared to wt IAV infection was specific for the tested TEs, as the ratio between cytosolic and nuclear localization was comparable between the two infections for other RNA species, such as MALAT1, GAPDH and the viral M segment (**Figs 3N-O, Fig EV3**). These data further indicate that IAV-induced TE-dsRNAs can reach the cytosol of infected cells, but that this phenomenon is particularly pronounced in the absence of the viral NS1 protein.

### IAV NS1 Protein Interacts with Infection-Induced TE-dsRNAs

Next, we wanted to address the possible mechanism by which NS1 expression may affect the localization of TE-dsRNAs. Given that we observed enhanced cytosolic localization of TE-dsRNAs during infection with IAV ΔNS1, that NS1 is known to bind non-specifically to dsRNA (Hatada & Fukuda, 1992), and that a substantial fraction of NS1 resides in the nucleus at early times post infection (**Fig EV4**)(Kerry *et al*, 2011), we hypothesized that NS1 might be able to interact with IAV-induced TE-dsRNAs as a mean to limit their transport into the cytosol. To this end, we immunoprecipitated transiently expressed V5-tagged wt NS1, or a well-characterized mutant NS1 (R38A/K41A) that lacks dsRNA binding activity (Wang *et al*, 1999), from HEK293T cells and incubated the bead-captured NS1s (or V5-tagged control, GST) with total RNA extracted from IAV-infected A549 cells (**Fig 4A**). Following this immunoprecipitation, protein extracts were assessed by western blot to validate equal immunoprecipitation of V5-tagged GST, wt NS1 and NS1-R38A/K41A (**Fig 4B**). RT-qPCR analysis of co-precipitated RNA revealed that (as expected) V5-tagged wt NS1, but not V5-tagged NS1-R38A/K41A could co-precipitate a fraction of the IAV M segment RNA (Hatada *et al*., 1992). Furthermore, several candidate TE-dsRNAs, including LTR13, MER72, MER21B, MLT1F and MLT2B3 were specifically co-precipitated with V5-tagged wt NS1, but not with V5-tagged NS1-R38A/K41A (**Fig 4C**). These data expand the range of dsRNA molecules that NS1 can form a complex with and suggest a plausible mechanism by which NS1 expression can limit cytosolic appearance of host TE-dsRNAs: interaction and sequestration in the nucleus.

**Figure 4:**
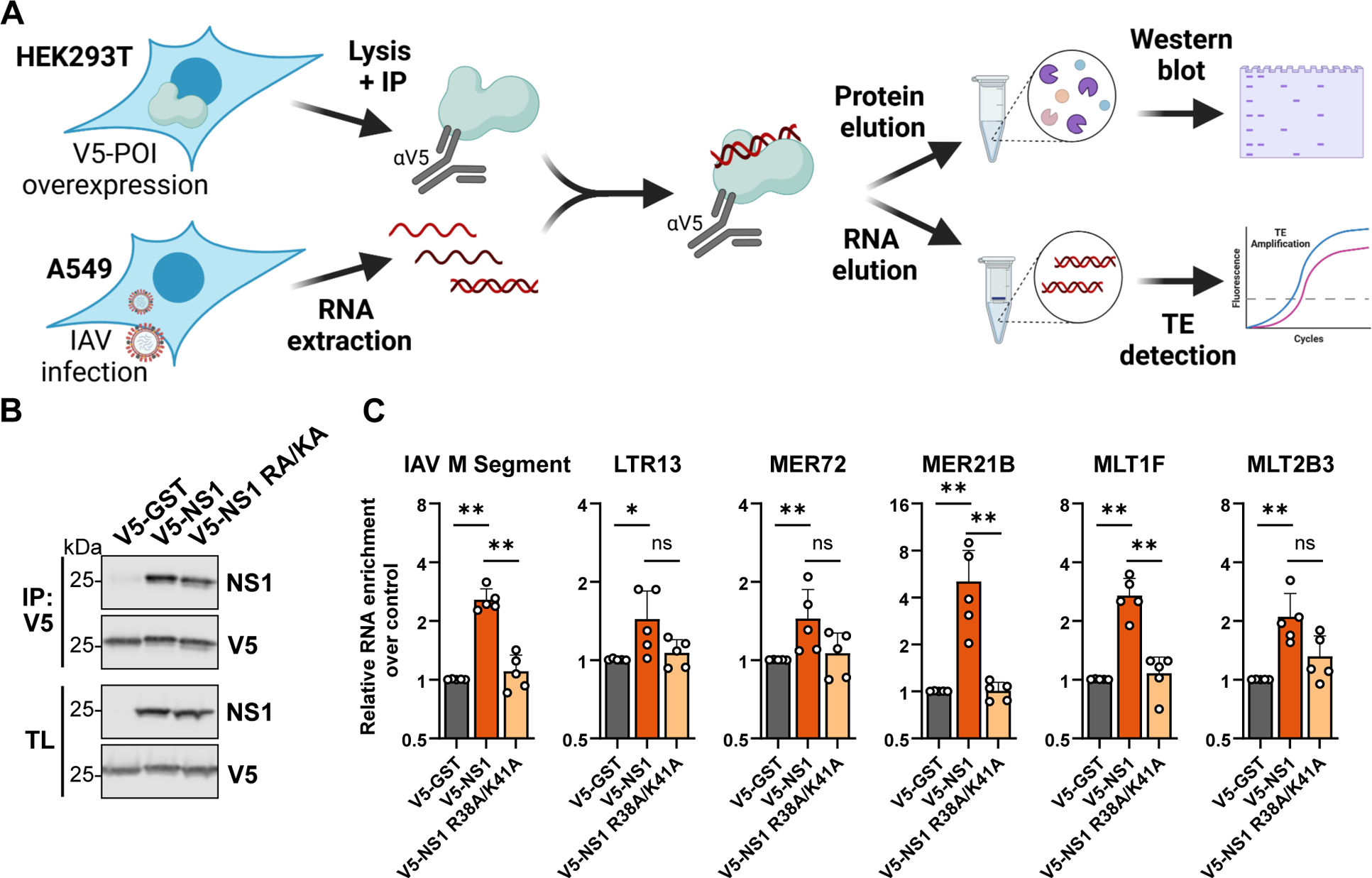
IAV NS1 Protein Interacts with Infection-Induced TE-dsRNAs. **A.** Experimental set-up for NS1 IP experiments: schematic representation of the experimental design aimed at delineating if NS1 interacts with de-repressed TEs. V5-tagged proteins of interest (POI) were immunoprecipitated from transfected HEK293T cells, and the immunoprecipitates were incubated with total RNA extracted from IAV-infected A549 cells [MOI 5 PFU/ml; 16 hpi]. Selected TEs were quantified by RT-qPCR. **B.** Western blot analysis of immunoprecipitated proteins: western blot analysis illustrating the successful immunoprecipitation of V5-tagged GST, wt NS1, or a dsRNA-binding deficient mutant NS1 (R38A/K41A; RA/KA) from transfected HEK293T cells. Total lysate (TL) served as a transfection control. Data are representative of n=5 independent experiments. **C.** RT-qPCR analysis of co-precipitating TEs: RT-qPCR analysis of selected TEs co-precipitated with wt NS1 or the dsRNA-binding deficient mutant NS1 (R38A/K41A; RA/KA). Enrichment is presented as fold change over the negative control (V5-GST). Bars represent mean values and SDs from n=5 independent experiments (each dot corresponds to one replicate). Statistical significance was determined by Mann-Whitney U test (*P ≤ 0.05; **P ≤ 0.01; ns, non-significant).

### IAV-Induced TE-dsRNAs can be Engaged by a Broad Range of Cytosolic Antiviral Sensors

We aimed to understand the possible fates of IAV-induced TE-dsRNAs that escape NS1 sequestration and that re-localize to the cytosol. Given that lack of NS1 expression leads to enhanced immunostimulatory activity in IAV-infected cells ((Yang *et al*, 2023) and our own data reported here), we focused on the potential recognition of TE-dsRNAs by host antiviral dsRNA sensors. Various cytosolic dsRNA-binding proteins have been implicated in the detection of endogenous dsRNAs such as Alu duplexes, LINE elements and ERVs, including the conventional viral sensors RIG-I (Ahmad *et al*, 2018; Du *et al*, 2023; Zhao *et al*, 2018) and melanoma differentiation-associated protein 5 (MDA5) (Ahmad *et al*., 2018; Chiappinelli *et al*., 2015; Roulois *et al*., 2015; Zhao *et al*., 2018), as well as protein kinase R (PKR)(Kim *et al*, 2018), Z-NA binding protein (ZBP1) (Devos *et al*, 2020; Jiao *et al*, 2020; Maelfait *et al*, 2017), Staufen1 (Ku *et al*, 2021), and adenosine deaminase acting on RNA 1 (ADAR1) (Chung *et al*, 2018). To evaluate the possible interaction of IAV-induced TE-dsRNAs with these RNA binding proteins, we therefore employed our previously described approach (**Fig 4A**) to ectopically express each V5-tagged sensor protein in HEK293T cells, followed by their immunoprecipitation and incubation with total RNA extracted from IAV-infected A549 cells. Following immunoprecipitation, enrichment of specific TE RNAs was assessed by RT-qPCR. Successful immunoprecipitation of the different dsRNA-binding proteins was verified by western blot analysis (**Fig 5A**). As expected, we observed specific enrichment of IAV M segment RNA with RIG-I (Rehwinkel *et al*, 2010) and ZBP1 (Zhang *et al*, 2020), but not with any of the other dsRNA-binding proteins (**Fig 5B**). Strikingly, our panel of candidate TE-dsRNAs (LTR13, MER72, MER21B, MLT1F, MLT2B3) exhibited a much broader reactivity to the dsRNA-binding protein panel: we observed that the TEs were precipitated not only by RIG-I, but also by MDA5, ZBP1, and PKR (**Fig 5B**). However, there was still a degree of specificity observed, as these TEs were not co-precipitated with either Staufen1 or ADAR1 (isoform p150). These data indicate that IAV-induced host TE-dsRNAs have the potential to be recognized by a range of cellular antiviral dsRNA-binding proteins should these components localize to the same cytosolic compartment. Given the breadth of immune and inflammatory pathways that can be activated by these different sensors (Hur, 2019), this observation highlights how multiple immunological consequences may result from IAV-induced TE-dsRNA de-repression, as well as the important role that NS1-mediated sequestration of TE-dsRNA likely plays in mitigating such effects.

**Figure 5:**
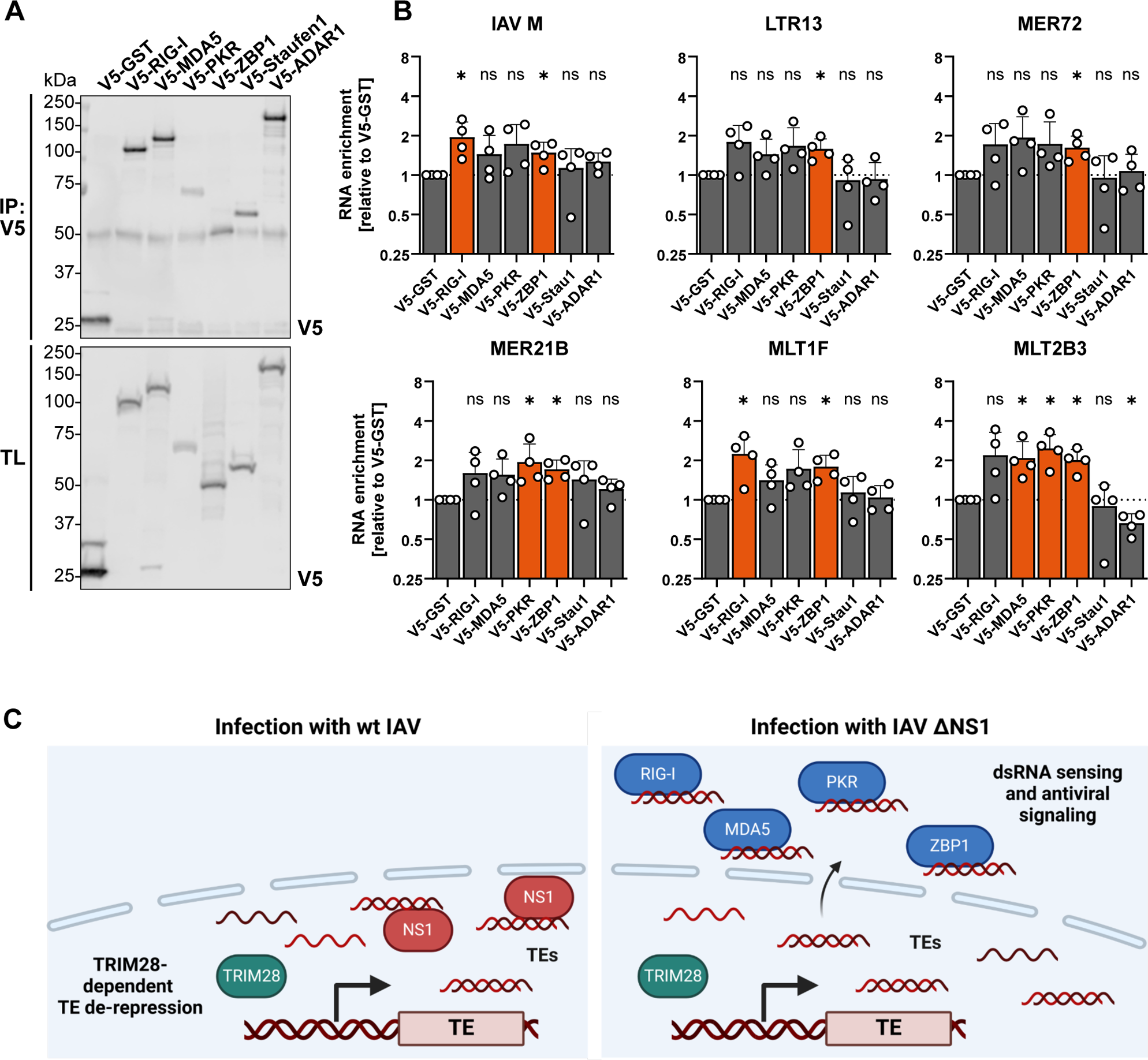
IAV-induced TE-dsRNAs can be Engaged by a Broad Range of Cytosolic Antiviral Sensors. **A.** Western blot of immunoprecipitated proteins: western blot analysis illustrating the successful immunoprecipitation of the indicated V5-tagged proteins from transfected HEK293T cells following experiments similar to Fig. 4A. Total lysate (TL) served as a transfection control. Data are representative of n=4 independent experiments. **B.** RT-qPCR analysis of co-precipitating TEs: V5-tagged proteins were immunoprecipitated from transfected HEK293T cells and subsequently incubated with total RNA extracted from IAV-infected A549 cells [MOI 5 PFU/ml; 16 hpi]. Selected TEs were quantified by RT-qPCR. Enrichment is illustrated as fold change over the negative control (V5-GST). Bars represent mean values and SDs from n=4 independent experiments (each dot corresponds to one replicate). Statistical significance was determined by one-sample t-test on log-transformed data (*P ≤ 0.05; ns, non-significant). Significantly enriched samples are highlighted in orange. **C.** Model of NS1-mediated inhibition of TE cytosolic translocation: schematic representation delineating the model of NS1’s role in hindering the translocation of de-repressed TEs into the cytosol where they are sensed by factors such as RIG-I, MDA5, PKR or ZBP1.

## Discussion

In this study, we used comprehensive RNA sequencing, computational analysis, and experimental approaches to characterize expression and localization of TEs in response to IAV infection. Our results demonstrate that IAV induces substantial genome-wide de-repression of TEs, and that TE RNAs can have a double-stranded nature. Furthermore, while these TE-dsRNAs are predominantly localized in the nucleus during infection with wt IAV, infection with a recombinant IAV lacking expression of the dsRNA-binding protein, NS1, results in their increased cytosolic presence (model in **Fig 5C**). A plausible mechanism by which NS1 expression prevents the re-localization of TE-dsRNAs from the nucleus to the cytosol was suggested by the interaction of NS1 with TE-dsRNAs. Lastly, we describe that TE-dsRNAs escaping NS1 sequestration can be engaged by a broad range of host antiviral sensor proteins with multiple immune and inflammatory functions. Our results therefore provide new insights into potential roles for these sensor proteins in IAV infection irrespective of their ability to directly recognize viral genomic material.

Previous studies have characterized TE expression in response to infection with different viruses and found that infection-induced genome-wide TE expression often occurs in the vicinity of antiviral response genes (Chen *et al*., 2023; Macchietto *et al*., 2020). Thus, de-repression of TEs might have *cis*-acting consequences on regulating the expression of these antiviral response genes (Chuong *et al*, 2017). However, our study adds an additional layer to virus-induced TE-mediated regulation of antiviral immunity: we analyzed the localization of TE transcripts and their potential to form dsRNA structures, hallmarks for successful detection by cytosolic host innate immune sensors. Our data therefore indicate that the widespread virus-induced upregulation of TE-dsRNAs might be an additional line of defense activated by the cell in response to virus infection, with these host-derived TE-dsRNAs potentially acting in a similar way to viral ‘pathogen-associated molecular patterns’ (PAMPs) to stimulate antiviral defenses. It is presently unclear why such an additional set of ‘host PAMPs’ may be generated and what role they would play over and above viral PAMPs. However, one could speculate that given many viruses have evolved measures to hide their own genetic material from host detection (e.g. by replicating in specialized bodies or protecting their vRNA 5ʹ ends through cap addition) (Markiewicz *et al*, 2021), as well as antagonistic mechanisms to target and inhibit specific sensing pathways (e.g. IAV NS1 targeting RIG-I) (Ayllon & García-Sastre, 2015), TE-dsRNA induction and broad activation of multiple sensors might be an effective cellular ‘mechanism of last resort’ to amplify innate and inflammatory immune responses.

Numerous dsRNA-binding proteins have been associated with detection of endogenous dsRNAs. For example, RIG-I and MDA5, the classical cytosolic sensors of viral dsRNAs, were also proposed to bind to ‘self’ LINE and Alu elements in the absence of infection during autoimmune conditions and in radiation-induced immune responses (Ahmad *et al*., 2018; Du *et al*., 2023; Zhao *et al*., 2018). MDA5 was further associated with sensing ERVs upregulated in response to treatment with DNA methyltransferase inhibitors, which block the DNA methylation important for epigenetic silencing of TEs (Ahmad *et al*., 2018; Chiappinelli *et al*., 2015; Roulois *et al*., 2015; Zhao *et al*., 2018). In uninfected cells, PKR can be activated by binding to dsRNAs formed by inverted Alu repeats (IRAlus) (Chung *et al*., 2018; Kim *et al*, 2014). Moreover, ZBP1, a protein that specifically binds to dsRNA in an alternative Z-conformation, was associated with binding viral and endogenous RNAs during viral infection or in the absence of SETDB1, resulting in induction of inflammation and cell death pathways (Devos *et al*., 2020; Jiao *et al*., 2020; Maelfait *et al*., 2017; Wang *et al*, 2020). Furthermore, the dsRNA-binding protein Staufen1 has been shown to stabilize endogenous retrovirus RNAs induced by DNA-methyltransferase inhibitors and thus transform cells into the viral mimicry state resulting in cell death (Ku *et al*., 2021). ADAR1, an enzyme converting adenosines to inosines, destabilizes dsRNA-structures formed by Alu elements to prevent their recognition as virus-derived dsRNAs (Chung *et al*., 2018). In this study, we observed that multiple dsRNA-binding proteins, including RIG-I, MDA5, PKR, and ZBP1, have the capacity to interact with de-repressed TEs during IAV infection, suggesting a potential defensive fallback within the cellular context. In scenarios where one of these proteins is directly counteracted by a virus, alternative candidates may therefore remain available to assume regulatory functions. Thus, we uncover the possibility that many different host sensors play previously unappreciated roles in virus infections irrespective of their direct recognition of viral genomic material.

Given that infection-triggered TE-dependent induction of an antiviral response (by viral mimicry) poses a powerful threat to the invading virus, it is conceivable that viruses may have developed antagonistic measures to counteract this. Indeed, we could show that IAV NS1 can interact with TE-dsRNAs, potentially sequestering them in the nucleus and preventing their detection by cytosolic sensors. Although NS1 is widely recognized as an antagonist of the host immune response on many levels, our recent findings unveil an additional dimension to its multifaceted repertoire and provide insights into a previously unappreciated possible host-derived dsRNA target for this viral protein. NS1 was initially described to bind to viral dsRNA formed during viral replication (Hatada & Fukuda, 1992; Hatada *et al*., 1992). However, there is still some controversy over whether the genomes of negative strand RNA viruses, such as IAV, do form aberrant dsRNAs during infection (Son *et al*., 2015; Weber *et al*., 2006). It is conceivable that other viruses may employ similar strategies to target potentially immunostimulatory host dsRNAs and prevent their recognition. For example, several viruses, such as vaccinia virus or Ebola virus, are also known to harbor dsRNA-binding proteins (E3L and VP35, respectively) (Cárdenas *et al*, 2006; Chang *et al*, 1992) that might elicit such a function. In contrast, other viruses might have developed distinct countermeasures, for example by using replication strategies to avoid TE de-repression or by directly targeting specific upstream or downstream mechanisms.

In conclusion, our study sheds light on the interactions between IAV and TEs within infected host cells (**Fig 5C**). We demonstrated that IAV NS1 can interact with de-repressed TEs induced during infection, likely sequestering most of them in the nucleus and thereby influencing their subcellular distribution and immunostimulatory potential. Furthermore, we investigated the fate of de-repressed TEs in the cytosol, revealing their recognition by a broad set of antiviral dsRNA sensors, including RIG-I, MDA5, ZBP1, and PKR. These findings underscore the multifaceted interplay between viral proteins, host cellular processes, and the regulation of TE expression during IAV infection, providing potential insights into the mechanisms underlying viral pathogenesis and host-activated immune or inflammatory responses.

## Methods

### Cells and Viruses

A549, HEK293T, and MDCK cells were cultured in Dulbecco’s modified Eagle’s medium (DMEM) (Life Technologies) supplemented with 10% (vol/vol) FCS, 100 units/ml penicillin, and 100 μg/ml streptomycin (Gibco Life Technologies). A549-ACE2/TMPRSS2 cells were a kind gift from Prof. Sam Wilson (University of Cambridge, UK) (Rihn *et al*, 2021). A549-ACE2/TMPRSS2 cells were used for the initial transcriptomics analysis, while A549 cells were used for all follow-up experiments. IAV strain A/WSN/1933 (WSN; H1N1) was propagated in MDCK cells. A recombinant WSN virus lacking NS1 expression (IAV ΔNS1) was kindly provided by Dr. Balaji Manicassamy (University of Iowa, USA) (Manicassamy *et al*, 2010) and was propagated in MDCK cells stably expressing the IAV NS1 protein (PR8 strain) (Hale *et al*, 2006) and the NPro protein from Bovine Viral Diarrhea Virus (Hilton *et al*, 2006). Following the initial RNA sequencing experiments, we discovered that the IAV ΔNS1 stock used was inadvertently contaminated with parainfluenza virus 5 (PIV5). A new non-contaminated IAV ΔNS1 stock was prepared, and small-scale experiments comparing TE expression in response to both the contaminated (S1) and non-contaminated (S2) IAV ΔNS1 stocks revealed identical TE responses (**Fig EV5**). We therefore conclude that the contamination did not grossly impact the large-scale RNA sequencing results as presented. All subsequent validation and follow-up experiments were performed with PCR-confirmed non-contaminated IAV ΔNS1 stocks.

### Virus Infection

Generally, cells were seeded at 6 × 10^5^ cells per well of a 6-well plate or 2.5 × 10^6^ cells per 10cm culture dish and infected the next day at the indicated multiplicity of infection (MOI). Virus inoculum was prepared in PBS supplemented with 0.3% BSA, 1 mM Ca^2+^/Mg^2+^, 100 units/ml penicillin, and 100 μg/ml streptomycin. Cells were incubated with the inoculum for 1 h, washed with PBS, and then overlaid with DMEM supplemented with 0.1% FBS, 0.3% BSA, 20 mM Hepes, 100 units/ml penicillin, and 100 μg/ml streptomycin. For the transcriptome analysis three independent replicates were performed.

### Plasmids

Mammalian expression vectors for pCAGGS-GST-V5, pLVX-PR8-NS1-wt-V5 and pLVX-PR8-NS1-R38A/K41A-V5 were described previously (Hale *et al*, 2008; Lopes *et al*, 2017). pcDNA3.1-V5-RIG-I, pcDNA3.1-V5-MDA5 and pcDNA3.1-V5-ADAR1 were generated with GeneArt (Thermo Fisher Scientific). ZBP1, PKR and Staufen1 were cloned into pcDNA3.1-V5 from pcDNA-3xFlag-ZBP1synth (generated with GeneArt, Thermo Fisher Scientific), pCAGGS-FLAG-PKR (kind gift from Adolfo García-Sastre, New York) and pCMV-Tag2B-Staufen1 (kind gift from Norbert Bannert, Berlin).

### Subcellular Fractionation

Cytosolic and nuclear fractions were isolated using a protocol adapted from Conrad *et al* (Conrad & Ørom, 2017), and total RNA was subsequently extracted from these fractions. Briefly, cells were seeded in 6-well plates and infected 24 h later for the indicated times. At the point of harvest, cells were detached by trypsinization, centrifuged at 1,200 rpm, and the resulting cell pellets were washed two times with ice-cold PBS. Cells were lysed for 5 minutes on ice in 100 μl cell lysis buffer (10 mM Tris pH 7.4, 150 mM NaCl, 0.15 % IGEPAL CA-630; sterile filtered through a Steritop-GP 0.22 μm filter unit and freshly supplemented with cOmplete™ Protease Inhibitor Cocktail (#11836170, Roche) and RNasin® Ribonuclease Inhibitor (#N2615, Promega; 1:1000)). Lysates were gently overlaid on top of 250 μl ice-cold sucrose buffer (10 mM Tris pH 7.4, 150 mM NaCl, 24 % sucrose; sterile filtered through a Steritop-GP 0.22 μm filter unit and freshly supplemented with RNasin® (1:1000)) in protein LoBind 1.5 ml tubes and centrifuged at 3500 × g for 10 min. The resulting supernatant containing the cytoplasmic fraction was cleared by centrifugation at 14,000 × g for 1 min in a new 1.5 ml microcentrifuge tube and the supernatant was collected. For RNA isolation from this fraction, 1 ml of Trizol (#15596026, Thermo Fisher Scientific) was added per 200 μl of cytosolic fraction. The pellet fraction containing cell nuclei was briefly rinsed with 1 ml ice-cold PBS-EDTA (1x PBS, 500 μM EDTA pH 8.0; freshly supplemented with RNasin® (1:1000)), followed by a 1-minute centrifugation at 3500 × g before gently removing the PBS-EDTA from the nuclear pellet. For RNA isolation from this fraction, nuclear pellets were directly lysed in 1 ml of Trizol reagent.

### RNA Extraction

Cells, lysates, or nuclei were directly lysed in 1 ml of Trizol (#15596026, Thermo Fisher Scientific) and incubated at room temperature (RT) for 10 minutes. 200 μl chloroform was added, samples were mixed by rigorous shaking, and centrifuged for 15 minutes at 13,000 rpm at 4°C. The aqueous phase was transferred to a new RNase-free tube containing 1.3 μl GlycoBlue™ Coprecipitant (#AM9515, Thermo Fisher Scientific), 1 volume of isopropanol (600 μl) was added, and after rigorous shaking samples were centrifuged for 15 minutes at 13,000 rpm at 4°C. The pellet containing RNA was washed twice with ice-cold 70% ethanol using centrifugation steps of 10 minutes at 13,000 rpm at 4°C. RNA was subsequently dissolved in nuclease-free water. RNA was used immediately for reverse transcription and the remaining RNA was stored at −80°C.

### RT-qPCR

Extracted RNA was reverse transcribed using the SuperScript™ IV First-Strand Synthesis System (#18091300, Thermo Fisher Scientific) with oligo(dT) primers and random hexamers according to the manufacturer’s instructions. RT-qPCR was then performed using PowerTrack™ SYBR Green Master Mix for qPCR (#A46109, Thermo Fisher Scientific) using specific forward and reverse primers listed in **Table EV1** in a 7300 Real-Time PCR System (Applied Biosystems). The relative gene expression was calculated with the ΔΔCt method (Livak & Schmittgen, 2001), using 18s-rRNA for normalization.

### Library Preparation and Sequencing

Library preparation and sequencing were performed by the Functional Genomics Center Zurich (FGCZ). Briefly, the quality of isolated RNA was determined with a Fragment Analyzer (Agilent, Santa Clara, California, USA). The TruSeq Stranded Total RNA Library Prep Gold (Illumina, Inc, California, USA) was used in subsequent steps. Briefly, total RNA samples (100-1000 ng) were depleted of ribosomal RNA and then reverse-transcribed into double-stranded cDNA. The cDNA samples were fragmented, end-repaired and adenylated before ligation of TruSeq adapters containing unique dual indices (UDI) for multiplexing. Fragments containing TruSeq adapters on both ends were selectively enriched with PCR. The quality and quantity of the enriched libraries were validated using a Fragment Analyzer (Agilent, Santa Clara, California, USA). The product was a smear with an average fragment size of approximately 260 bp. The libraries were normalized to 10nM in Tris-Cl 10 mM, pH8.5 with 0.1% Tween 20. A Novaseq 6000 (Illumina, Inc, California, USA) was used for cluster generation and sequencing according to a standard protocol. Sequencing was paired-end at 2×150 bp.

### Sequencing Data Analysis

Reads were trimmed and base corrected using *fastp-0.23.2* (Chen *et al*, 2018) and reads shorter than 35 nucleotides were excluded. The quality of the trimmed reads was checked using FastQC-0.12.1 (Andrews, 2010). Trimmed reads were then aligned to the hg38 human genome using *STAR-2.7.10b* (Dobin *et al*, 2013), keeping only reads with a single best alignment and a maximum number of 3 mismatches. A custom *Rust* (Matsakis, 2014) program was written to count the number of potentially double-stranded reads. When provided with TE coordinates and the aligned reads, the program counts how many reads from each strand map to each TE locus and overlap with reads from the opposite strand. This results in two counts of reads per locus: the number of reads from the forward strand that overlap with reads from the reverse strand, and *vice versa*. The minimum of these counts was taken as an estimate of the number of double-stranded reads, assuming that the remaining reads are single-stranded. The read counts for non-TE genes were obtained using *featureCounts* from the *subread-2.0.1* package (Liao *et al*, 2014). An *R-4.3.1* script (R Core Team, 2023) was written to calculate the differential expression. Differential expression analysis of TE loci was performed using *DESeq2* (Love *et al*, 2014) using size factors calculated from non-TE read counts. Data transformations and visualizations were performed using *dplyr* (Wickham H, 2023) and *Rsamtools* (Morgan, 2023).

### dsRNA Immunoprecipitation

A549 cells seeded in 10cm culture dishes were infected with wt IAV or IAV ΔNS1 [MOI 5 PFU/cell; 16h] and total RNA was extracted by Trizol-chloroform extraction as described above. For each condition, 2 μl anti-dsRNA [9D5] (#Ab00458-1.1, Absolute Antibody) or mouse monoclonal anti-Flag M2 (#F1804, Sigma Aldrich), were coupled to 30 μl Protein G Sepharose™ beads (#P3296, Millipore) by 2 h incubation at 4°C with rotation. Washed beads were incubated with 30 μg RNA in 300 μl PBS + 0.1% Triton X-100 (freshly supplemented with1 μl RNasin®) for another 2 hours at 4°C with rotation, followed by 4 washes with PBS + 0.1% Triton X-100 (freshly supplemented with RNasin® (1:1000)). RNA was subsequently eluted and extracted using Trizol.

### Co-immunoprecipitation of RNA with Proteins

HEK293T cells (∼6 × 10^5^ cells per 6-well) were transfected with 1-3 μg of the indicated constructs using FuGENE® HD Transfection Reagent (#E2311, Promega) according to the manufacturer’s instructions. After approximately 24 hours, cells were washed once with PBS and lysed for 15 minutes on ice in lysis buffer (50mM Tris-HCL, pH 7.4, 300mM NaCl, 1mM EDTA, 1% Triton X-100) supplemented with cOmplete™ Protease Inhibitor Cocktail (#11836170, Roche). Lysates were then cleared by centrifugation for 15 minutes at 14,000 rpm at 4°C and subjected to immunoprecipitation with anti-V5 mouse monoclonal antibody (#MCA1360, Bio-Rad) overnight at 4°C with rotation in DNA/RNA LoBind tubes. Subsequently, 30 μl of Protein G Sepharose™ beads (#P3296, Millipore) were added and incubated for 2 hours at 4°C with rotation. The beads were washed twice with lysis buffer. The bead-captured immunoprecipitates were then incubated with 30 μg total extracted RNA from IAV [MOI 5 PFU/cell; 16h]-infected A549 cells in 300 μl PBS + 0.1% Triton X-100, freshly supplemented with 1 μl RNasin®, for 2 hours at 4°C with rotation. Following this, beads were washed 4 times with PBS + 0.1% Triton X-100 (freshly supplemented with RNasin® (1:1000)). Bound RNAs were eluted and extracted using Trizol as described above.

### Western Blot Analysis

Cell lysates were generated by lysis in 2× urea disruption buffer (6 M urea, 2 M β-mercaptoethanol, 4% SDS, bromophenol blue) followed by sonication to shear nucleic acids. Proteins were separated by SDS-PAGE, transferred to nitrocellulose membranes (Amersham), and detected using the following primary and secondary antibodies: mouse monoclonal GAPDH antibody (0411)(#sc-47724, Santa Cruz), rabbit polyclonal Histone H3 antibody (#ab1791, Abcam), rabbit polyclonal Influenza A virus NS1 antibody (#GTX125990, GeneTex), mouse monoclonal V5 antibody (#MCA1360, Bio-Rad), IRDye 680CW goat anti-mouse IgG (#926–68070, Li-Cor), and IRDye 800CW goat anti-rabbit IgG (#926–32211, Li-Cor).

### Statistical Analysis

Statistical analyses were generally performed using GraphPad Prism 7 software. For RT-qPCR data, ΔCt values were analyzed. The statistical tests used and P values for significance are stated in the figure legends. Gene ontology enrichment analyses were performed on DEGs (log_2_FC > 2 and Padj < 0.1) using DAVID (Huang da *et al*, 2009). Dot plot visualization of the Top 10 enriched GO terms for each condition (P Value < 0.001) was generated in R version 4.3.2 (R Core Team, 2023). TE classes were manually annotated according to Repbase (Bao *et al*, 2015).

## Acknowledgements

We thank Sam Wilson for A549-ACE2/TMPRSS2 cells, Balaji Manicassamy for the IAV ΔNS1 virus, and Adolfo García-Sastre and Norbert Bannert for plasmids. The authors gratefully acknowledge the Functional Genomics Center Zurich (FGCZ) of University of Zurich and ETH Zurich, for assisting with the RNA-seq experiment. Overview panels in figures 1A, 3A, 4A and 5C were created with BioRender.com. The research leading to these results received funding from the Swiss National Science Foundation (grant 310030_214957 to BGH) and the University of Zurich (Forschungskredit grant FK-22-051 to ML). The funders had no role in study design, data collection, data interpretation, or the decision to submit the work for publication.

## Author contributions

Conceptualization: ML and BGH; Methodology and Investigation: ML, LC, and GL; Writing and Visualization: ML and BGH; Supervision: RK and BGH; Funding Acquisition and Project Administration: BGH.

## Supporting information

Expanded View Figures PDF

Table EV1

Dataset EV1

Dataset EV2

## Data availability

The datasets produced in this study have been deposited to the European Nucleotide Archive (ENA) with the dataset identifier PRJEB75711. Code is available upon request.

