## Supplementary material for "Influenza A Virus NS1 Limits Recognition of Double-Stranded Transposable Elements by Cytosolic RNA Sensors": EV Figures

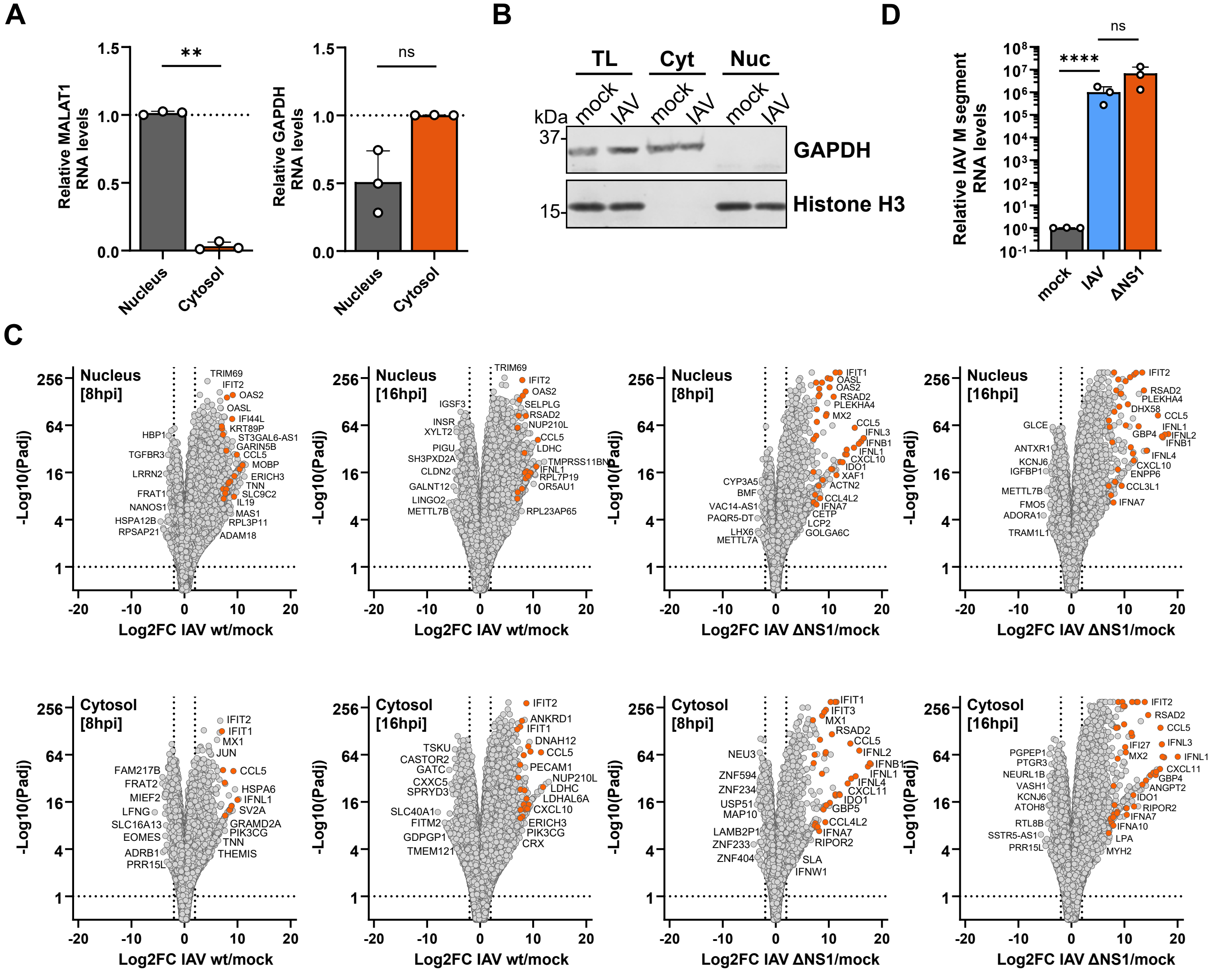


**Figure EV1: Fractionation Quality Controls and Subcellular Host Transcriptome Analysis During IAV Infection.**

**A**. Subcellular fraction purity assessment: RT-qPCR analysis was conducted on RNA samples from similar experiments to those shown in Fig 1 to evaluate the purity of subcellular fractions using specific markers. GAPDH was utilized as a cytosolic fraction marker, and MALAT1 served as a nuclear fraction marker. Bars represent mean values and SDs from n=3 independent experiments (each dot corresponds to one replicate). Significance was determined by unpaired t-test on log-transformed data (**$P \leq 0.01$; ns, non-significant).

**B**. Subcellular fraction purity assessment by western blot: western blot analysis was employed on protein samples from similar experiments to those shown in Fig 1 to assess subcellular fraction purity, probing for the specific marker proteins GAPDH (cytosolic fraction) and Histone H3 (nuclear fraction). Data are representative of n=3 independent experiments.

**C**. Differential gene expression analysis in subcellular fractions: volcano plots depicting the differential gene expression in A549-ACE2/TMPRSS2 cells upon infection with wt IAV or IAV ΔNS1 [MOI = 5 PFU/cell] at the indicated times and in the indicated subcellular fractions. Genes with statistically significant differential expression (Padj < 0.1) and a $log_2FC > 2$ are considered differentially expressed genes (DEGs). Cytokines, chemokines, and interferon-stimulated genes (ISGs) with a $log_2FC > 7$ are highlighted in orange, emphasizing genes with substantial expression changes. Selected gene names are shown.

**Figure EV1: Fractionation Quality Controls and Subcellular Host Transcriptome Analysis During IAV Infection. (Continued)**

**D**. Assessment of infection consistency between wt IAV and IAV ΔNS1: A549 cells were infected with wt IAV or IAV ΔNS1 [MOI = 5 PFU/cell] for 16 h. RT-qPCR analysis for IAV M segment was performed to confirm similar infection levels between wt IAV and IAV ΔNS1. Significance was determined by ordinary one-way ANOVA on log-transformed data (****$P \leq 0.0001$; ns, non-significant).

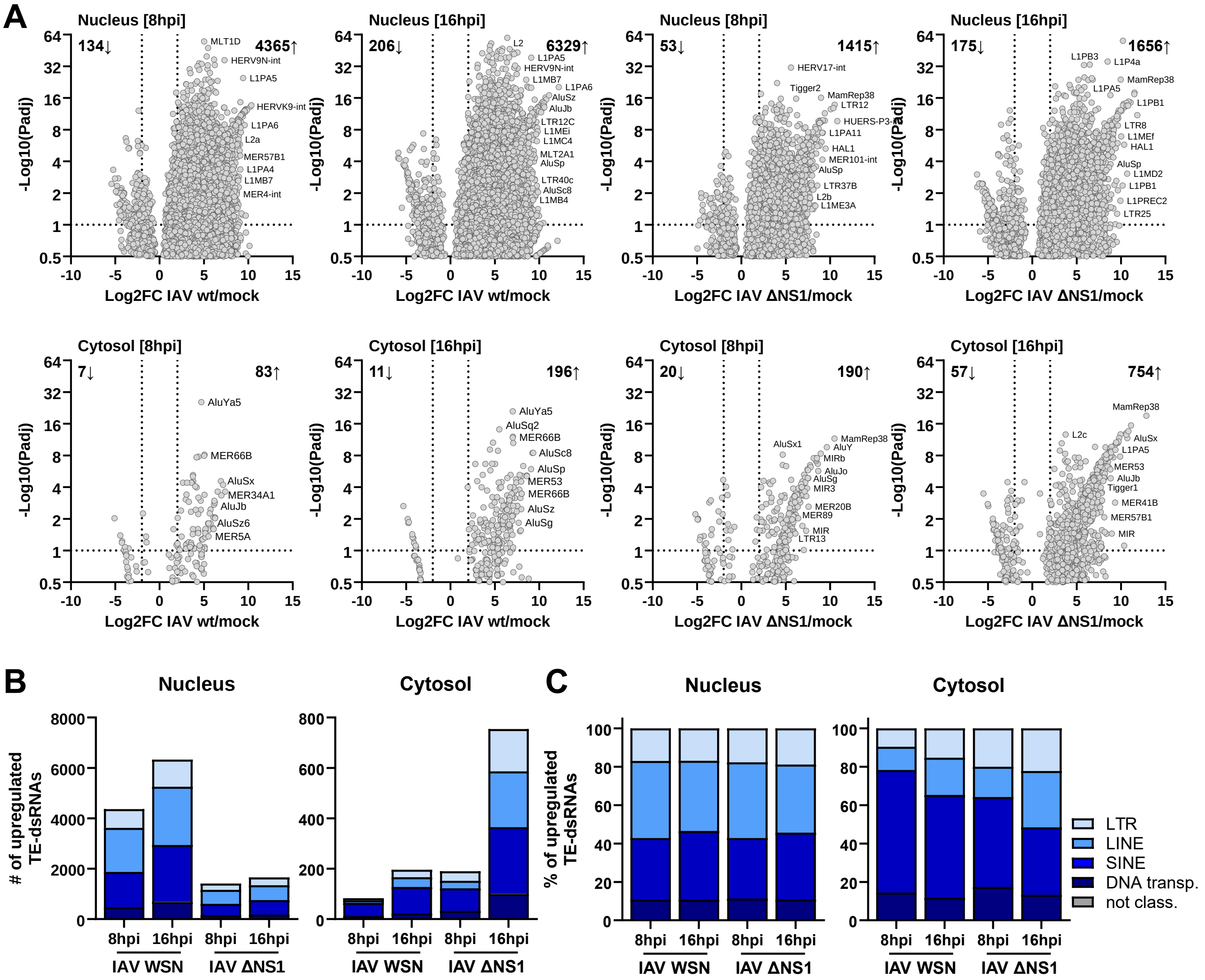


**Figure EV2: Differential Expression and Classification of Double-Stranded Transposable Elements in Subcellular Fractions During IAV Infection.**

**A**. Differential expression of double-stranded TEs in subcellular fractions: volcano plots illustrating the differential expression of double-stranded TEs in A549-ACE2/TMPRSS2 cells following infection with wt IAV or IAV ΔNS1 [MOI = 5 PFU/cell] in the specified subcellular fractions. Genes meeting criteria for statistical significance (Padj < 0.1) and a $log_2FC > 2$ are considered differentially expressed transposable elements (DE-TEs). Selected elements are labeled.

**B/C**. Subclasses of significantly upregulated double-stranded TEs in the cytosolic fraction: (**B**) absolute and (**C**) relative numbers of TE subclasses significantly upregulated during wt IAV or IAV ΔNS1 infections at the times indicated are shown for LTR (long-terminal repeat retrotransposon), LINE (long interspersed nuclear element), SINE (short interspersed nuclear element) and DNA transposon elements.

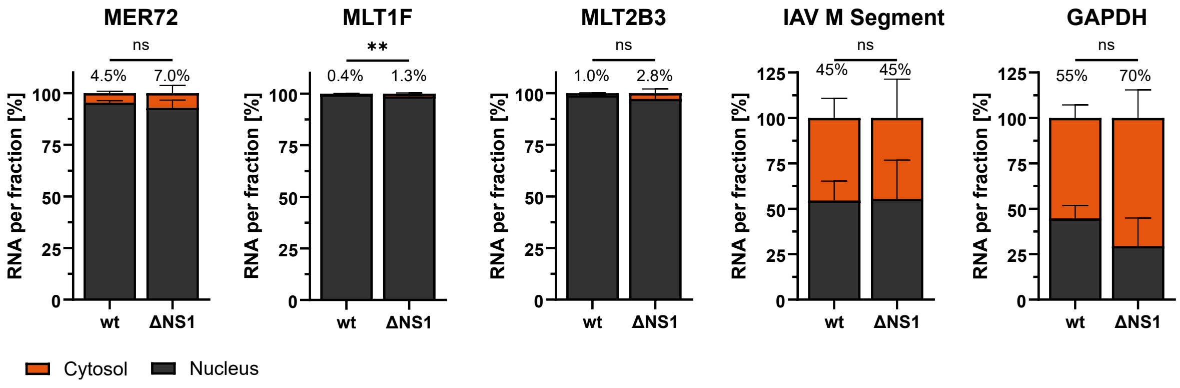


**Figure EV3: IAV-Induced Transposable Elements and their Increased Cytosolic Abundance in the Absence of NS1.**

TE expression ratios: expression of selected TEs was quantified by RT-qPCR (see 3C-J), and presented as the percentage of transcript per fraction. Bars represent mean values and SDs from n=4 independent experiments. The percentages annotated atop the bars specifically denote the cytosolic fractions. Significance was determined by unpaired t test (**P ≤ 0.01; ns, non-significant).

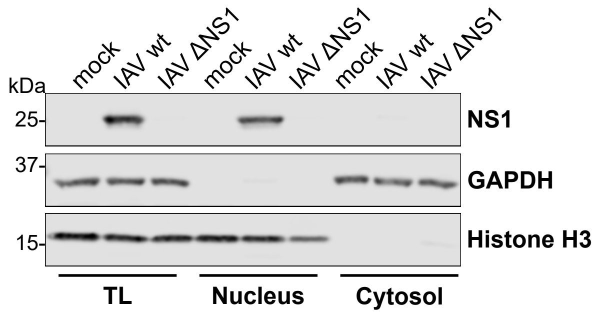


**Figure EV4: Predominant Nuclear Localization of IAV NS1 During Infection.**

Western blot analysis of IAV NS1 expression in subcellular fractions of A549 cells infected with wt IAV, IAV ΔNS1, or mock [MOI = 5 PFU/cell] for 16 h. The purity of subcellular fractions was also assessed by probing for the specific marker proteins GAPDH (cytosolic fraction) and Histone H3 (nuclear fraction). Data are representative of n=3 independent experiments.

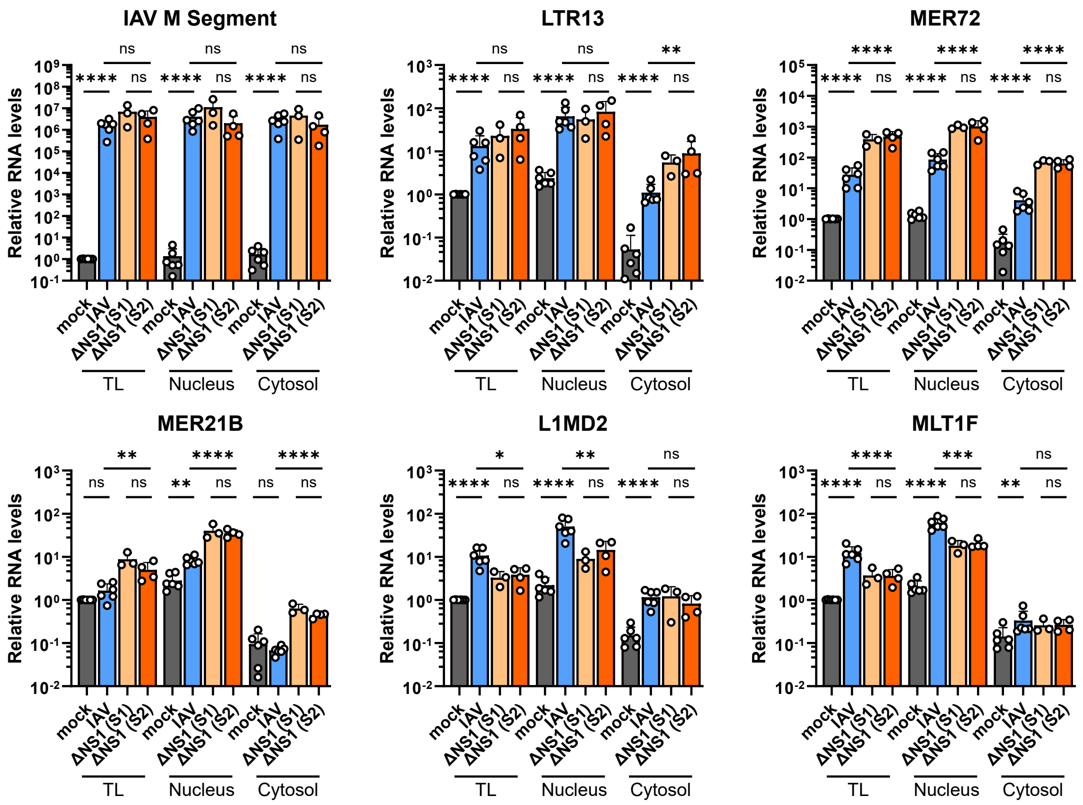


**Figure EV5: Validation of Differential TE Expression with Different IAV ΔNS1 Stocks.**

RT-qPCR analysis to validate the differential expression of selected TEs in subcellular fractions following mock, wt IAV, or IAV ΔNS1 infections [MOI = 5 PFU/cell] in A549 cells. The initial IAV ΔNS1 stock (S1) used for the large-scale sequencing experiment was subsequently discovered to be accidentally contaminated with parainfluenza virus 5 (PIV5). A new contamination-free IAV ΔNS1 stock (S2) was prepared and did not exhibit any differences in induction of TE expression or localization as compared to the original S1 contaminated stock. The data for mock, wt IAV, and IAV ΔNS1 (S2) are also shown in Fig. 3. Significance was determined by ordinary one-way ANOVA with Šídák's multiple comparisons test on log-transformed data (*$P \le 0.05$; **$P \le 0.01$; ***$P \le 0.001$; ****$P \le 0.0001$; ns, non-significant).
