## Supplementary material for "Influenza A Virus NS1 Limits Recognition of Double-Stranded Transposable Elements by Cytosolic RNA Sensors": EV Table 1

**Table EV1: Primer pairs used for RT-qPCR**

| **Name** | **Fw/Rv** | **Sequence** |
| --- | --- | --- |
| **GAPDH** | Fw | CTGGCGTCTTCACCACCATGG |
| **GAPDH** | Rv | CATCACGCCACAGTTTCCCGG |
| **MALAT1** | Fw | GAAGGAAGGAGCGCTAACGA |
| **MALAT1** | Rv | TACCAACCACTCGCTTTCCC |
| **IAV M (WHO M30F2)** | Fw | ATGAGYCTTYTAACCGAGGTCGAAACG |
| **IAV M (WHO M264R3)** | Rv | TGGACAAANCGTCTACGCTGCAG |
| **18s-rRNA** | Fw | GGCCCTGTAATTGGAATGACTC |
| **18s-rRNA** | Rv | CCAAGATCCAACTACGAGCTT |
| **LTR13** | Fw | AAGCTGGCCCACAGTTATCC |
| **LTR13** | Rv | AGGTGCACAGAGTGGAACAG |
| **MER72** | Fw | TGATCAGCCTTCCCTCCTGA |
| **MER72** | Rv | AGGACATAGAGGCACCCTGT |
| **L1MD2** | Fw | CTTCCCAGTTGTACAGGCGT |
| **L1MD2** | Rv | GCGTGATGGATGGTCGATCT |
| **MLT1F** | Fw | GGCATATGGCTTCCAAGGCT |
| **MLT1F** | Rv | ACATGGCCCTCTCCATAGGT |
| **MLT2B3** | Fw | GTTCTTGGCCTTTGGCCTTG |
| **MLT2B3** | Rv | CTGTAAGCTGGAGACCGTGG |
